# Direction and distance dependency of reaching movements of lower limb

**DOI:** 10.1101/2023.03.14.530922

**Authors:** Kyosuke Oku, Shinsuke Tanaka, Noriyuki Kida

## Abstract

Efficient movement of the body is an important part of our daily lives, which we perform unconsciously. Such efficient movements have been investigated using reaching movements in the upper limb, and whose kinematics reportedly vary depending on the movement direction. However, the reaching movements of the lower limbs are not well understood. Therefore, we aimed to examine changes in the kinematics of the lower limb reaching movements to determine the mechanism of skilled motor ability in terms of direction and distance. Sixteen adults (10 males) were requested to reach targets projected on the floor in seven directions and at three distances, for a total of 21 points. We found that the reaching time slowed down for the contralateral side (right foot to left-sided target), and was caused by a slower start of the toe movement. To identify the cause of this delay, we analyzed the onset of movement at each joint and found that movement to the contralateral side began closer to the torso than movement to the ipsilateral side. The time-to-peak velocity was also calculated; the motion required to reach the target in the shortest time varied depending on the direction and distance. Acceleration phases that were not observed in the upper limb velocity profile were observed in the lower limb velocity profiles. These results suggest that movement kinematics vary with direction and distance, resulting in slower reaching time on the contralateral side.

## Introduction

The central nervous system (CNS) unconsciously executes efficient movements daily. To ensure appropriate movement output, various factors should be considered, including the surrounding environment and an individual’s physical condition. For example, it is necessary to change direction and calculate the target distance when moving. Efficient reaching movements in the upper limb for direction and distance has been investigated [1, 2]. In particular, the peak velocity of the fingertips and the reaching time depends on the movement direction [3]. Furthermore, different strategies that reach the target the fastest are used [4]. These actions are thought to be the brain optimizing its control of the body to move faster and more efficiently [5]. However, this change in the reaching movement kinematics with respect to direction and distance is limited to the upper limb. Because reaching movements of the lower limbs involve a shift in the center of pressure (CoP), its kinematics are expected to differ from those of the upper limbs. Therefore, understanding the movement of the lower limbs would lead to a better understanding of the whole-body movement.

Reaching movements of the lower limbs for multiple directions and distances involve the CoP shift, and hence, are more complex than those of the upper limbs. When the lower limbs reach the contralateral side, the CoP shifts in the direction of movement, which may influence the accuracy and efficacy of the movement. The CoP reportedly shifts before the toes lift off the ground when taking the first step [6-8]. Moreover, the CoP transitions change with speed and direction [9]. However, to date, no study has examined the movement kinematics in terms of direction and distance and the CoP shifts. Therefore, the interaction between multiple distances and directions and motions involving CoP shifts should be examined.

The upper limb movements at various distances [1] and directions [10-12] have been characterized. A study examining the interaction between five distances and five directions did not find any correlation at accelerating or maximum speeds [4]. In contrast to the upper limbs, the lower limbs must support the body while controlling the legs. This role is thought to be a burden on the performance of lower limbs depending on the direction and distance. Therefore, we performed a lower limb reaching test to determine the interaction between direction and distance.

Motion analysis of 3D data is a powerful and promising approach for characterizing the movement linked to the CoP shift. Most previous studies have used force plate data [7, 8]; no studies have used 3D data till date. Additionally, it is important to understand the mechanisms by which the terminal motor unit moves the trunk. Here, we aimed to evaluate the reaching movement of the lower limb using 3D motion data and determine how the motor skills varied with direction and distance. This study sheds light on the movement kinematics of the lower limb in association with CoP transition.

## Methods

### Participants

Sixteen healthy adults (10 males and 6 females), aged 23.0 ± 2.55 years (mean ± SD), participated in this study. All participants were right-footed, as assessed using the Waterloo Footedness Questionnaire [13]. The participants were selected based on the criteria that they had normal vision and no experience playing soccer at a competitive level. We selected a sample size (n = 16) based on previous studies [14-19]. Prior to the experiments, its purpose and procedure were explained to the participants, and a written informed consent was obtained. This study was approved by the Ethics Committee of the Graduate School of Human and Environmental Studies, Kyoto University and was conducted in accordance with the Declaration of Helsinki.

### Experimental setup and apparatus

A schematic of the experiment is presented in Fig. 1. The participants were made to stand upright on a wooden platform (180 cm wide and 90 cm deep) placed in a dim room. A projector (H6530BD, Acer) placed approximately 200 cm above and 180 cm in front of the participant’s right foot projected the visual targets on the floor. The visual targets were white circles with a diameter of 10 cm that were drawn using a custom software written in Visual Basic 2017 (Microsoft Visual Studio, Microsoft) and run on Windows 10 via VGA outputs. A total of 21 visual targets in seven directions (22.5, 45.0, 67.5, 90.0, 112.5, 135.0, and 157.5°) and at three distances (25.0, 50.0, 75.0 cm) from the great toe of the right foot. The farthest distance from the visual target was set based on the approximate criterion that normal adults could reach with a single movement [20-22].

**Fig. 1.**
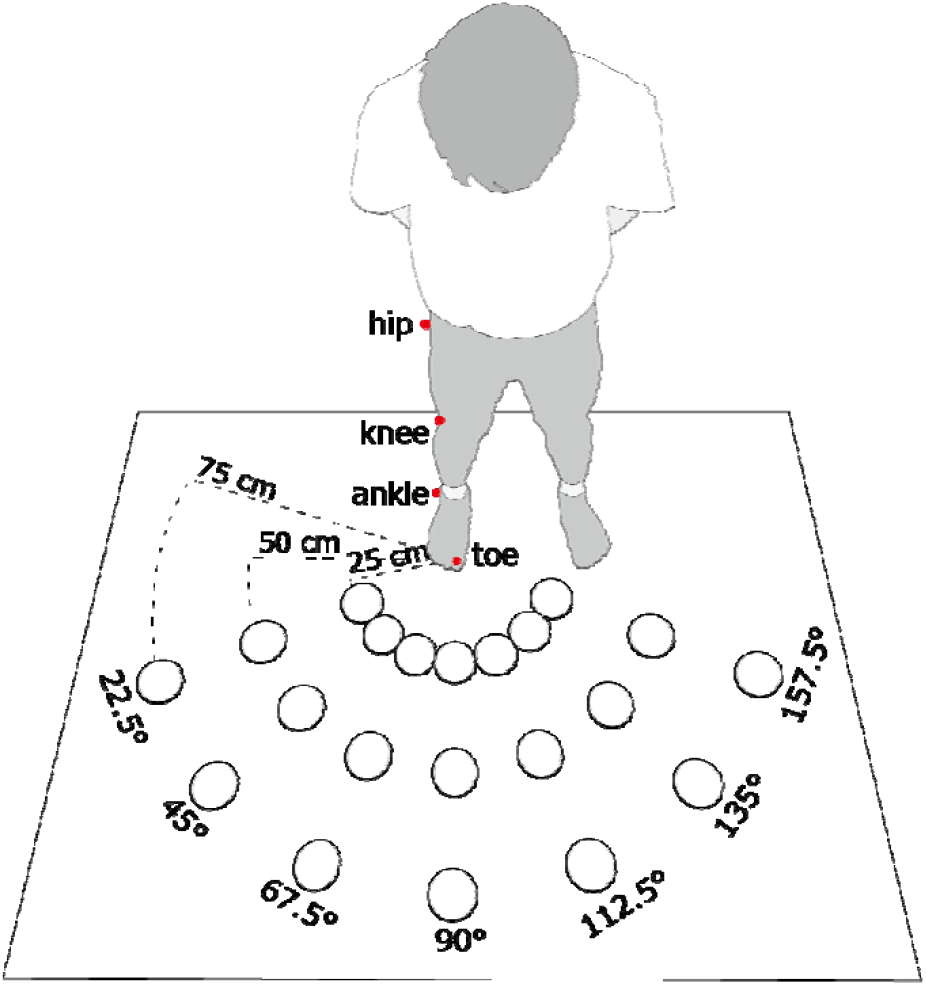
Overview of the experimental setup. It depicts the distances and directions of the projected visual stimuli and the location of the infrared markers.

Kinematic data were obtained using a motion-capture system (OptiTrack, NeuralPoint). Reflective markers of 4 mm diameter were placed on the tip of the right second toe, while those with a diameter of 16 mm were placed on the surface of the right ankle (lateral malleolus), knee (lateral femoral condyle), and hip (greater trochanter). The spatial locations of the four reflective markers were captured using six infrared cameras (OptiTrack Flex 3) with a temporal resolution of 100 Hz. A software-controlled infrared LED (tip diameter of 5 mm and wavelength of 850 nm) was placed on the floor and synchronously illuminated with the visual targets to enable detection of the visual stimulation onset using the motion-capture system.

### Procedure

First, the participants performed a motor task in which they reached the target with their right foot. The participants stood upright with their heels on the platform and stabilized their posture before starting the task. One of the 21 visual targets (seven directions and three distances) was presented to the participant with a 1–4 s random delay after using a beep to cue the onset. As soon as the participant visually perceived the target, he/she tried to reach it with his/her right foot as per the following instruction: “reach for the target with the toe as quickly and accurately as possible.” Upon reaching the target, the participants were asked to keep the toe on the target for 2 s to verify termination of one trial. Trials were consecutively conducted for all 21 locations with a 5-s interval. Subsequently, this set of 21 trials was repeated 4 times (84 trials in total), with a 2-min interval to prevent fatigue. The participants practiced the reaching task 10 times prior to the actual session.

### Data Analysis

The time series data of 3-dimensional location of the reflective markers were preprocessed by a second order Butterworth low-pass filter with a cutoff frequency at 10 Hz. The positional data were differentiated in time by 3-point differential algorism to obtain velocity on each moment for each point. The velocity in three axes in each frame for each point were synthesized to induce the instantaneous velocity in 3-dimentional space. The onset and offset of the movement were defined by each criterion in each point. For the toe, the first of the 5 consecutive frames in which the velocity exceeded 30 cm/s was defined as the onset while the last of 5 consecutive frames in which the velocity fell below 30 cm/s was defined as offset of the movement. This threshold was set to 10 % of the speed of the fastest movement among the 21 points in the average of all participants. In the same manner, the criteria for onset and offset for each point were set as 28.0, 18.0, 9.6 cm/s for ankle, knee and hip, respectively.

To evaluate the kinematics of lower limb, we calculated the following indices; movement initiation (MI) for each point (MI_toe_, MI_ank_, MI_kne_ and MI_hip_ for toe, ankle, knee and hip, respectively) defined as duration from onset of target presentation to that of movement; movement time (MT) which was defined as duration from onset to termination of toe movement: task duration (TD) which was defined as sum of MI_toe_ and MT, indicating duration from onset of target presentation to termination of toe movement; time to peak velocity (TPV) which was defined as the duration from onset of target presentation to the appearance of maximum velocity for each point (TPV_toe_, TPV_ank_, TPV_kne_ and TPV_hip_ for toe, ankle, knee and hip, respectively).

Furthermore, we conducted a correlation analysis between each factor (TD, MI of each point, MT, TPV of each point) and direction to examine how movement change with direction. To examine how movement and cognition change with direction and distance, the MI of each point was analyzed using three-factor analysis of variance (3×7×4 ANOVA); distance, direction, and point (toe, ankle, knee, and hip). If a two-way interaction effect was detected, it was applied to a two-factor analysis of variance of direction and point for each of the three distances (7×4 ANOVA was performed three times). If a single interaction effect was detected, multiple comparisons of Bonferroni-corrected were performed to examine the differences in MI between points at each distance. For TPV of each point, Two-factors analysis of variance of direction and distance was also conducted (7×3 ANOVA was performed four times). If a single interaction effect or main effect was detected, multiple comparisons of Bonferroni-corrected were performed to examine how TPV differed with distance and direction for each point.

## Result

The velocity profile from initiation to completion is shown in Fig. 2. The velocity profiles of the toe and ankle were largely similar, depicting a single peak which increased with the distance moved. However, there were two peaks in the knee and hip as movement time increased; a large peak preceded a small peak in the knee, while the small peak preceded the large peak in the hip. In both cases, the trend was more prominent as the distance increased.

**Fig. 2.**
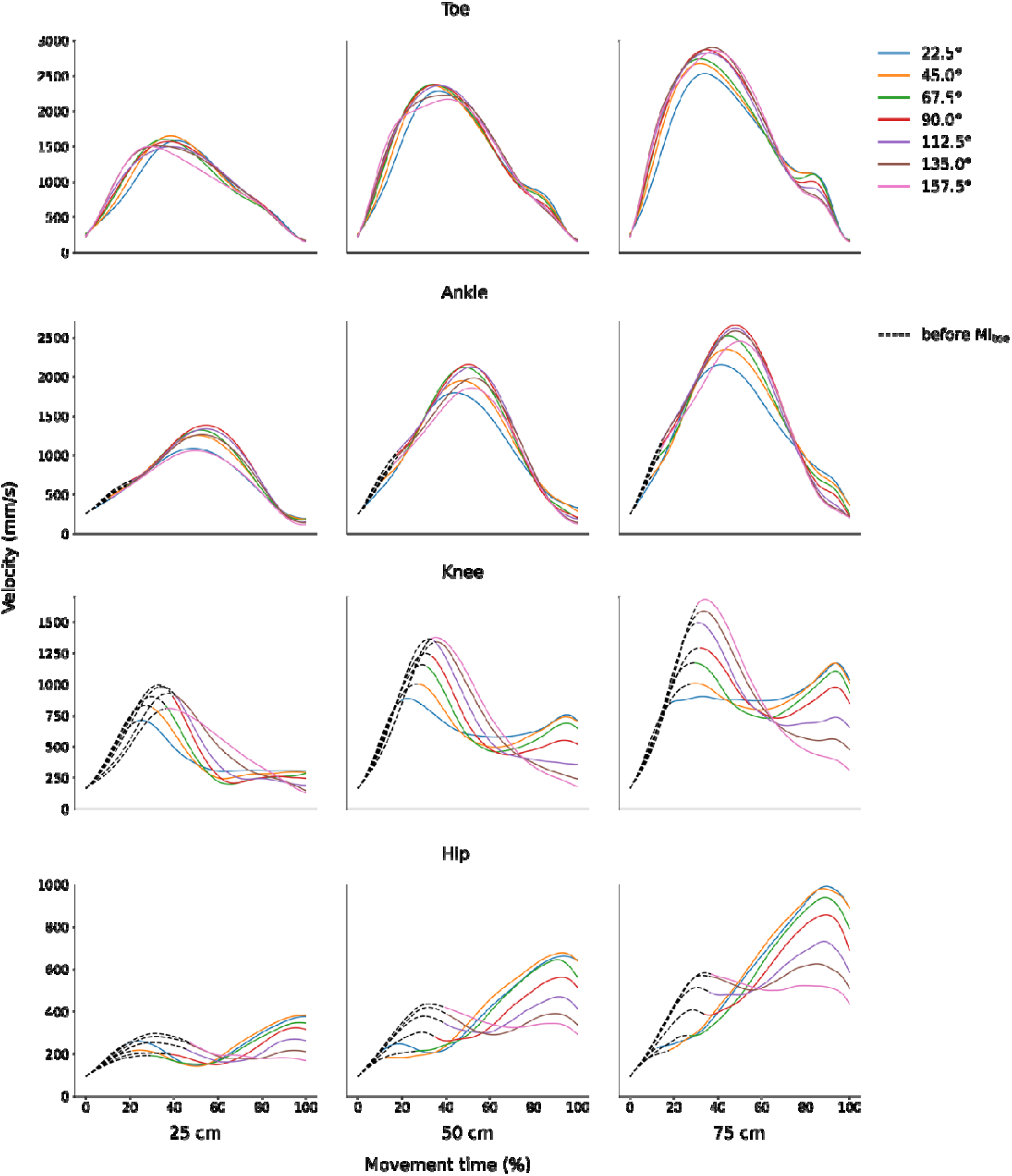
Velocity profile of each joint in relation to distance. The dotted line shows the change in velocity before initiation of toe movement.

The average values of the task duration (TD), movement initiation (MI), and movement time (MT) are shown in Fig. 3. TD increased with distance and direction angle, particularly up to a reaching distance of 50 cm (Fig. 3. left panel). To determine more details about body movements, we divided TD into MI of toe (MI_toe_) and MT; the average values are shown in Fig. 3 (right panel). MT tended to decrease with the increase in direction angle, whereas MI_toe_ appeared to increase with the increase in direction up to 90°. Thus, variation in TD can be largely explained by changes in MI_toe_ rather than those in MT.

**Fig. 3.**
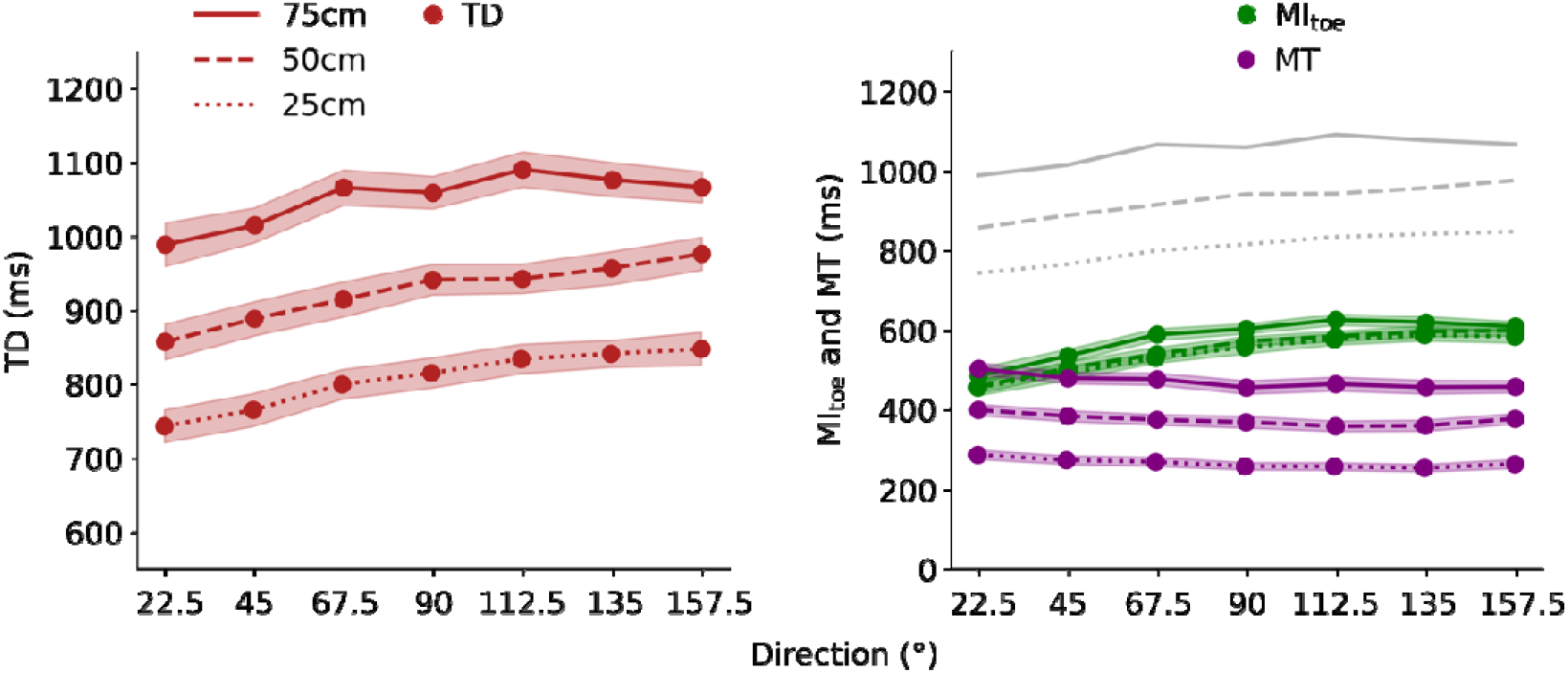
Left panel: TD is the duration from visual stimulus presentation to movement termination. Right panel: MI_toe_ is the duration from visual stimulus presentation to the movement initiation of the toe in actual motion; it represents the movement initiation in this study. MT is the duration from movement initiation to completion. The lightly colored bands in both panels indicate SE (Standard error). TD is shown in gray in the right panel for reference. TD, task duration; MI_toe_, movement initiation of the toe; MT, movement time.

The average MI values for each joint, which provide insight into the entire limb movement, are shown in Fig. 4. The MI for the toe and ankle similarly increased with increase in direction angle. However, this was not the case for hip MI. Direction dependency of MI_hip_ was biphasic; MI increased with increase in direction angle up to approximately 45°, and thereafter decreased with further increase in direction angle. This trend was particularly prominent at a reaching distance of 50 cm (Fig. 4. center panel).

**Fig. 4.**
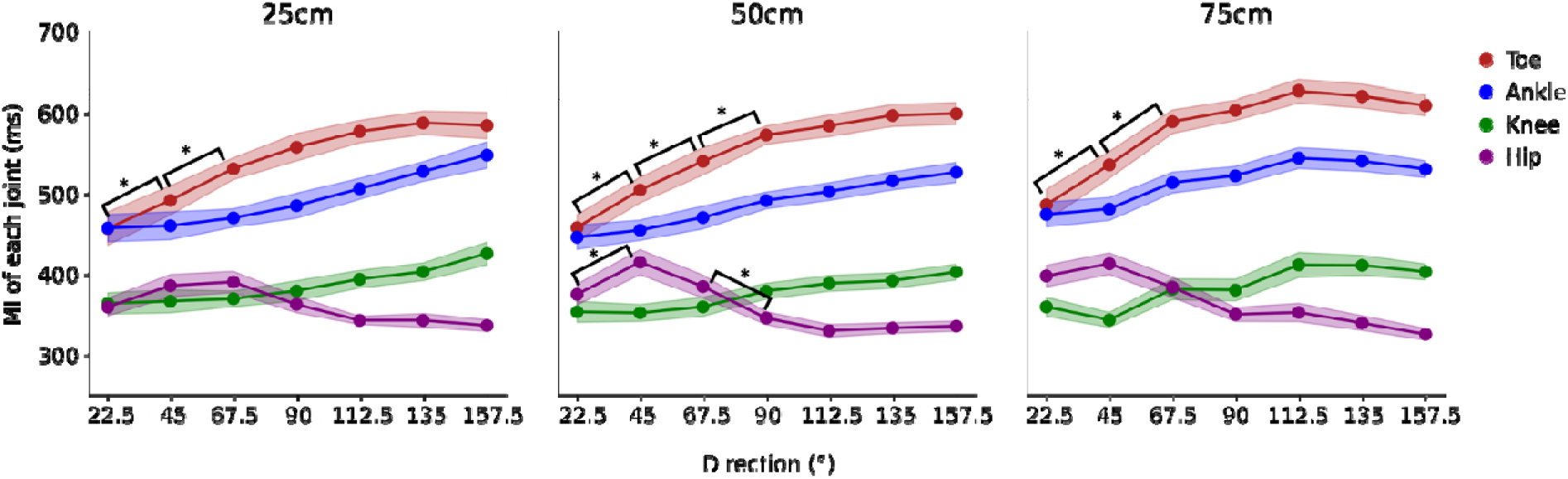
Movement initiation of each joint. The line color represents the joint, and the line the distance. The lightly colored bands indicate SE (Standard error). The significant difference between each joint at 0.5% level is shown.

The average values of the time-to-peak velocity (TPV) at each direction angle are shown in Fig. 5. TPV of the hip was higher than those of other joints, and increased with the increase in direction angle up to 90°; thereafter, it decreased. The TPV of the knee linearly increased with the increase in direction angle at a distance of 25 cm. In contrast, it decreased with the increase in direction angle up to approximately 90°; thereafter it increased. Thus, the TVP of the knee seems to converge to roughly the same value, irrespective of the distance, at higher direction angles (most contralateral side). The TPV of the ankle and toe largely increased with an increase in direction angle.

**Fig. 5.**
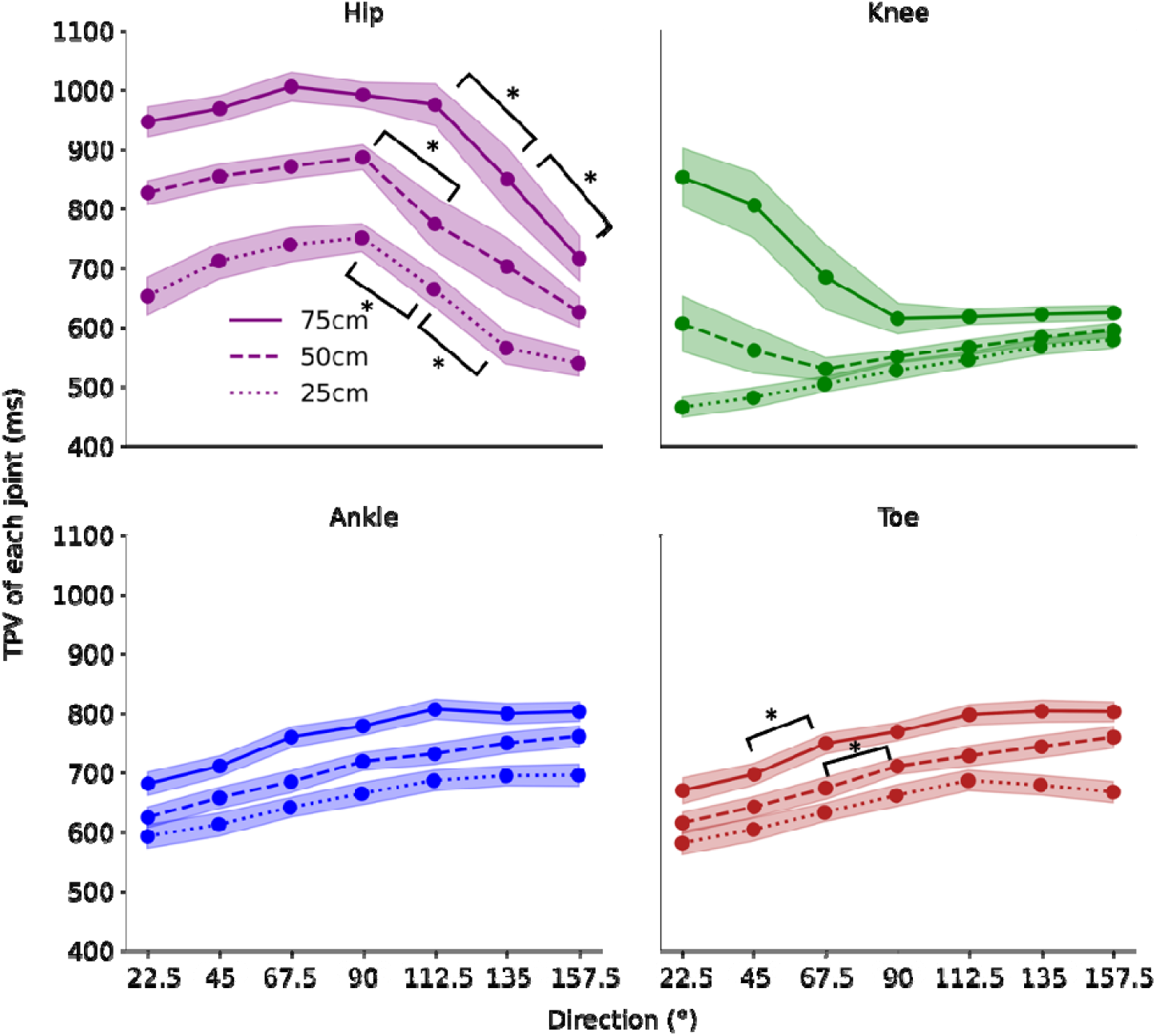
Time-to-peak velocity (TPV) in the hip, knee, ankle and toe. The lightly colored bands indicate the standard error (SE). Significant difference between each direction angle at 0.5% level is shown.

To understand the direction and distance dependency of the reaching ability, we analyzed the correlation between each kinematic parameter and the direction angle for all distances (25, 50, and 75 cm); the results are summarized in Table 1. There was a significant positive correlation between the TD and direction angle for all distances. The MI_toe_ also demonstrated a positive correlation with the direction angles. In contrast, a statistically significant negative correlation was noted for MT at distances of 25 and 75 cm, with a tendency of negative correlation at 50 cm.

**Table 1.**
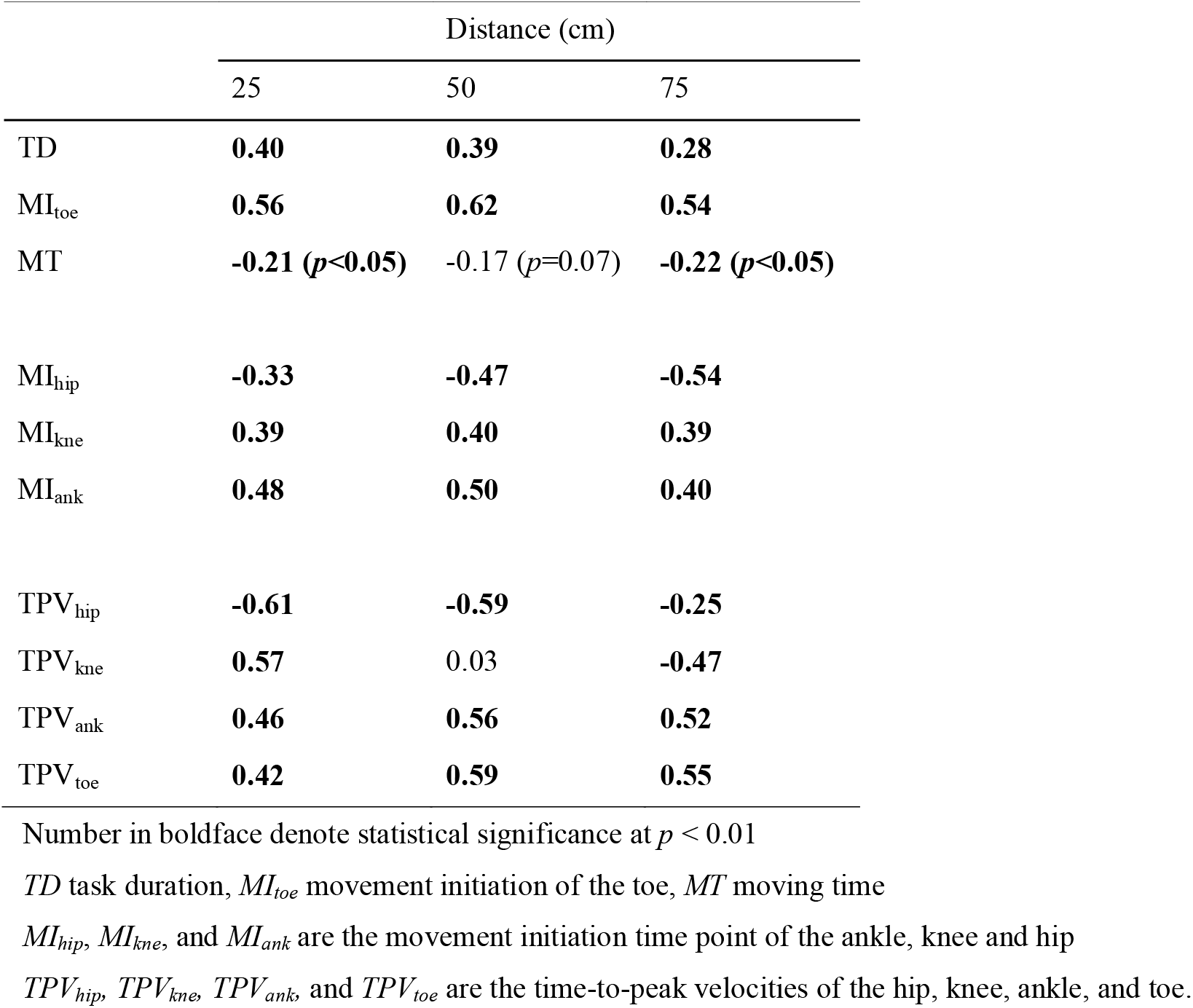
Correlation between the direction angle and each kinematic parameter (*r*)

|  | Distance (cm) |  |  |
| --- | --- | --- | --- |
|  | 25 | 50 | 75 |
| TD | **0.40** | **0.39** | **0.28** |
| $MI_{toe}$ | **0.56** | **0.62** | **0.54** |
| MT | **-0.21 ($p<0.05$)** | -0.17 ($p=0.07$) | **-0.22 ($p<0.05$)** |
| $MI_{hip}$ | **-0.33** | **-0.47** | **-0.54** |
| $MI_{kne}$ | **0.39** | **0.40** | **0.39** |
| $MI_{ank}$ | **0.48** | **0.50** | **0.40** |
| $TPV_{hip}$ | **-0.61** | **-0.59** | **-0.25** |
| $TPV_{kne}$ | **0.57** | 0.03 | **-0.47** |
| $TPV_{ank}$ | **0.46** | **0.56** | **0.52** |
| $TPV_{toe}$ | **0.42** | **0.59** | **0.55** |

Number in boldface denote statistical significance at $p < 0.01$
$TD$ task duration, $MI_{toe}$ movement initiation of the toe, $MT$ moving time
$MI_{hip}$ , $MI_{kne}$ , and $MI_{ank}$ are the movement initiation time point of the ankle, knee and hip
$TPV_{hip}$ , $TPV_{kne}$ , $TPV_{ank}$ and $TPV_{toe}$ are the time-to-peak velocities of the hip, knee, ankle, and toe.

The correlation between the direction angle and MI of each joint is summarized in Table 1. A significant positive correlation between the direction angle and toe and knee was confirmed for all distances; however, this was not seen in the hip (Table 1). MI_hip_ exhibited a negative correlation with the direction angle. A positive correlation between the MI and direction angle indicated that the initiation movement was delayed. The correlation coefficient appeared to be higher for the joints furthest from the trunk: MI_toe_ > MI_ank_ > MI_kne_ >MI_hip_. This suggested that the initial movement of the joint farthest from the trunk was more strongly influenced by the direction in which it is to be moved.

The results of the correlation analysis between the direction angle and TPV of each joint support the above observations (Table 1). TPV_hip_ showed a consistently negative correlation with direction angles at all distances; however, the coefficient was lower for longer distances (Fig. 4). The correlation of TPV_kne_ was completely opposite at 25 and 75 cm (Fig. 4); a positive correlation at 25 cm, no correlation at 50 cm, and a negative correlation at 70 cm with increasing direction angle. TPV_ank_ and TPV_toe_ exhibited a positive correlation with increase in direction angle at all distances, reflecting the direction dependency of the TPV (Fig. 4).

To determine the statistical significance of these correlations, we conducted a three-factor analysis of variance (ANOVA) for MI (3×7×4) using distance, direction, and joint (toe, ankle, knee, and hip). We found a significant two-way interaction effect (*F* [36, 540] = 5.69, *p* < 0.01, *η*^*2*^= 0.003), simple interaction effect between the joint and direction angle (*F* [18, 270] = 125.31, *p* < 0.01, *η*^*2*^ =0.068), and main joint effect (*F* [3, 45] = 391.40, *p* < 0.01, *η*^*2*^ = 0.64). Multiple comparisons of Bonferroni-corrected MI_toe_ data revealed a significant difference between 22.5° and 45° and between 45° and 67.5° at all distances. There was a significant difference in MI_hip_ duration between 22.5° and 45° and 67.5° and 90° at 50 cm. There was a significant difference between MI_toe_ and MI_ank_ at all three distances and direction angles > 22.5° (*p* > 0.05).

A two-factor ANOVA of direction and distance was applied for TPV (7×3 ANOVA was conducted four times). There was a significant simple interaction effect between distance and direction angle (*F* [12, 180] = 2.53, *p* < 0.01, *η*^*2*^ = 0.009) and main effect of distance (*F* [2, 30] = 99.73, *p* < 0.01, *η*^*2*^ = 0.230) and direction angle (*F* [6, 90] = 71.95, *p* < 0.01, *η*^*2*^ = 0.227) in the TPV_toe_. Multiple comparisons of the Bonferroni-corrected data revealed significant differences between 67.5° and 90° at 50 cm and between 45° and 67.5° at 75 cm (*p* < 0.05).

No significant simple interaction effect was noted for the TPV_ank_ between distance and direction angle (*F* [12, 180] = 1.35, *p* > 0.05, *η*^*2*^ = 0.004); there was a significant main effect of distance (*F* [2, 30] = 81.22, *p* < 0.01, *η*^*2*^= 0.226) and direction angle (*F* [6, 90] = 69.96, *p* < 0.01, *η*^*2*^ = 0.218). Multiple comparisons of the Bonferroni-corrected values revealed no significant difference between the adjacent direction angles (*p* > 0.05).

We found a significant simple interaction effect between distance and direction angle (*F* [12, 180] = 15.39, *p* < 0.01, *η*^*2*^ = 0.150) and a significant main effect of distance (*F* [2, 30] = 59.05, *p* < 0.01, *η*^*2*^ = 0.245) and direction angle (*F* [6, 90] = 3.52, *p* < 0.01, *η*^*2*^ = 0.032) in the TPV_kne_. Subsequent multiple comparisons of Bonferroni-corrected data revealed no significant differences between the adjacent direction angles (*p* > 0.05).

A significant simple interaction effect between distance and direction angle was observed for TPV_hip_ (*F* [12, 180] = 1.19, *p* < 0.05, *η*^*2*^ = 0.012). Additionally, there was a significant main effect of distance (*F* (2, 30) = 111.25, *p* < 0.01,*η*^*2*^ = 0.342) and direction angle (*F* [6, 90] = 42.30, *p* < 0.01, *η*^*2*^ = 0.220). Multiple comparisons of Bonferroni-corrected data confirmed a significant difference between 90° and 112.5°at 25 cm and 50 cm and between 112.5° and 135° at 75 cm (*p* < 0.05).

## Discussion

The TD distinctively increased with the increase in direction angle, indicating that movement to the contralateral side took more time than movement to the ipsilateral side (Fig. 3, left panel). This was largely due to higher MI_toe_ that could not be compensated for by the decrease in MT with increase in direction angle (Fig. 3, right panel). Furthermore, the correlation coefficient of MI_toe_ with TD was much higher than that of MT, indicating that the variations in TD were explained by changes in MI_toe_ rather than by those in MT. Thereafter, we analyzed the MI of each joint to determine why the MI_toe_ changed. The MI_toe_ and MT showed highly positive and negative correlations with the direction angle, respectively. The MI_toe_ increased with increase in direction angle predominated the decrease in MT on the contralateral side. This resulted in an overall increase in TD with increase in direction angle (gray lines in Fig. 3, right panel).

The MI of each joint suggested that the movement differs between the ipsilateral and contralateral sides. MI_toe_ and MI_ank_ were almost comparable for ipsilateral movement at 22.5°; however, the MI_ank_ was lower than MI_toe_ at the contralateral side beyond 45° (Fig. 4). The MI_hip_ was higher than MI_kne_ in the ipsilateral direction < 67.5°; this relationship was reversed for movements beyond 90°. This observation shows that the hip moved faster, followed by the knee, ankle, and toe, as the direction increased to the contralateral side. These results imply that the kicking motion on the contralateral side starts from the part of the body closest to the trunk. This is especially true for movement to the farthest contralateral direction, where the action was primed by the hip movement. Directional dependence of the movements is thought to be related to gait initiation. Gait initiation involves a CoP transition before the toes leave the ground. There are several reports on the relationship between various factors and CoP transition in preparation for walking [6-8]. The CoP shift reportedly differs depending on the direction [9]. Our study results are consistent with the view that movement to the contralateral side starts from the body part closest to the trunk. Thus, the movement preparation before toe-off differs between movement toward the ipsilateral and contralateral sides, which may consequently modulate the MI_toe_.

We also found differences in the PTV depending on the direction, distance, and joint (Fig. 5), which were similar to those obtained for the reaching movements of the upper limbs. TPV reportedly differs between movement in the horizontal direction [23], and between prehension movements of the upper limbs [12]. Similarly, TPV also differs in vertical up-and-down movements [5, 24]. The present study found more complex relationships between the joints and distances (Table 1). Overall, the hip showed a negative correlation with direction throughout, while the ankle and toe showed a consistent positive correlation. The trend of the knee was biphasic, which was similar to that of the ankle and toe for close distances (25 cm) and similar to that of the hip for far distances (75 cm). This implies that the part of the limb farthest from the trunk, the ankle and toe, moved consistently irrespective of the reaching distances and direction. The knee moved in a direction similar to the ankle or toe for close distances (25 cm) and similarly to the hip for far distances (70 cm). The movement to the farthest contralateral direction was probably primed by the movement of the hip, followed by that of the knee, ankle, and toe. Therefore, it seems plausible that knee movement integrates the action of the limb parts closest and farthest from the trunk and plays a pivotal role in reaching the object on the contralateral side. These findings may be explained by the aforementioned CoP shift [9]. However, TPV is thought to be more complex. We tested lower-limb reaching movements in this study and found that strategies for faster reaching may be complex and differ from those of the upper limb depending on distance, direction, and joint.

We observed two characteristics of note in the velocity profiles (Fig. 1). First, the hip velocity profile was bimodal. This was particularly evident on movements to the contralateral side and may have been caused by CoP transition. The first peak of the hip appeared before MI_toe_, which reached the contralateral side first and required movement transmission from the trunk to the toe. This bimodality was evident in our results that MI_hip_ was faster than MI_kne_ on the contralateral side and reinforced the notion of movement transmission (Fig. 5).

Second, the toe velocity profile differed to some extent from the bell shape described in previous studies. The upper limb reaching movements show a bell-shaped velocity profile [1, 25]. In our study, there was another shoulder peak just before the end of the movement, particularly at far distances. This may be due to the transmission of the second peaks of the knee or hip, which is a novel characteristic suggesting a connection of the joint parts in reaching movements. These characteristics discovered in this study are novel and unique to the lower limb.

## Limitations and future prospects

The current study investigated movements in a stationary environment; therefore, our results cannot be reproduced in a dynamic situation. In addition, we focused on distances and time from a stationary state; however, in the future, we would like to investigate interpersonal distances in a dynamic situation. In doing so, we hope to determine more advanced human functions related to distance and time.

## Conclusion

The current study provides the novel insight that the reaching movements of the lower limb may change with direction and distance. Kinematic data demonstrated that movement preparation varies with direction, which changes the timing of toe-off. This characteristic is specific to the lower limb. Overall, our findings highlight the involvement of joint transmission mechanisms in the lower limb.

## Acknowledgments

We would like to thank Professor Keisuke Kushiro for his beneficial advice. This study was supported by a grant-in-aid for scientific research (K.K.).

